# Cycledash: a web application for the interactive analysis and exploration of variants

**DOI:** 10.1101/125153

**Authors:** Isaac Hodes, Dan Vanderkam, Tavi Nathanson, B. Arman Aksoy, Jaclyn Perrone, Jeff Hammerbacher

## Abstract

As genomics begins to influence clinical care, validating the result of a somatic variant calling pipeline has become increasingly important. Cycledash is a web application which facilitates the validation and analysis of somatic variants, bringing together and streamlining existing tools and workflows for examining and verifying the existence of variants in patient samples.

**Availability:** The source code is freely available at https://github.com/hammerlab/cycledash under the Apache 2 License. A public demo (*username* cycledash *password* cycledash123) is available at http://cycledash.hammerlab.org. The web application is tested to work in recent version of Firefox, Safari, and Chrome, and the server applications will run on Linux and OS X systems.

## 1 Introduction

Identifying and understanding mutations in a patient’s DNA is becoming more vital to the treatment of cancer. Clinical variant calling pipelines can include manual quality control, some of which take significant amounts of time. These include examining the variant in the context of the short DNA reads in its vicinity (the pileup) using tools such as the Integrated Genomics Viewer (IGV); annotating the variants with inferred information such as their effect, the transcript they would appear in, and expression levels (using, for instance, Varcode); and filtering out low-depth variants, variants which appear in dbSNP, or variants with a very low frequency (Robinson et al., 2011; Rubinsteyn et al., 2016). All of these require complex decisions which are either manually carried out, or automated in a way that belies their underlying uncertainty.

In practice, VCF files are threaded through as series of applications, for example, Microsoft Excel, the Integrated Genomics Viewer, and Jupyter notebooks, which must be individually understood, each introducing a new set of possible errors. Cycledash was built to ameliorate these problems by unifying existing tools and workflows within an easy-to-use web application.

## 2 Implementation

Cycledash consists of a browser-based client and a server, as well as a worker queue for executing long-running jobs.

The server is written in Python using the Flask web framework. The server primarily exposes a RESTful API that the client queries for data, and which our variant calling pipeline can POST results to via HTTP. The server is also responsible for orchestrating the worker queue so that variants can be annotated. The exposed API is extensively tested using the nosetest framework. We use the PostgreSQL database to store our variants and related information, and RabbitMQ as our worker queue.

The client is written in JavaScript, using the React.js framework to coordinate user-interface updates in a predictable manner. The client can query the web API via AJAX to allow the user to responsively view very large variant files, as well as to examine variant calls in the context of their pileups. We use pileup.js (Vanderkam et al., 2016) to visualize BAM files and variant calls within them, displaying an IGV-like viewer within the table of variants.

**Figure 1:**
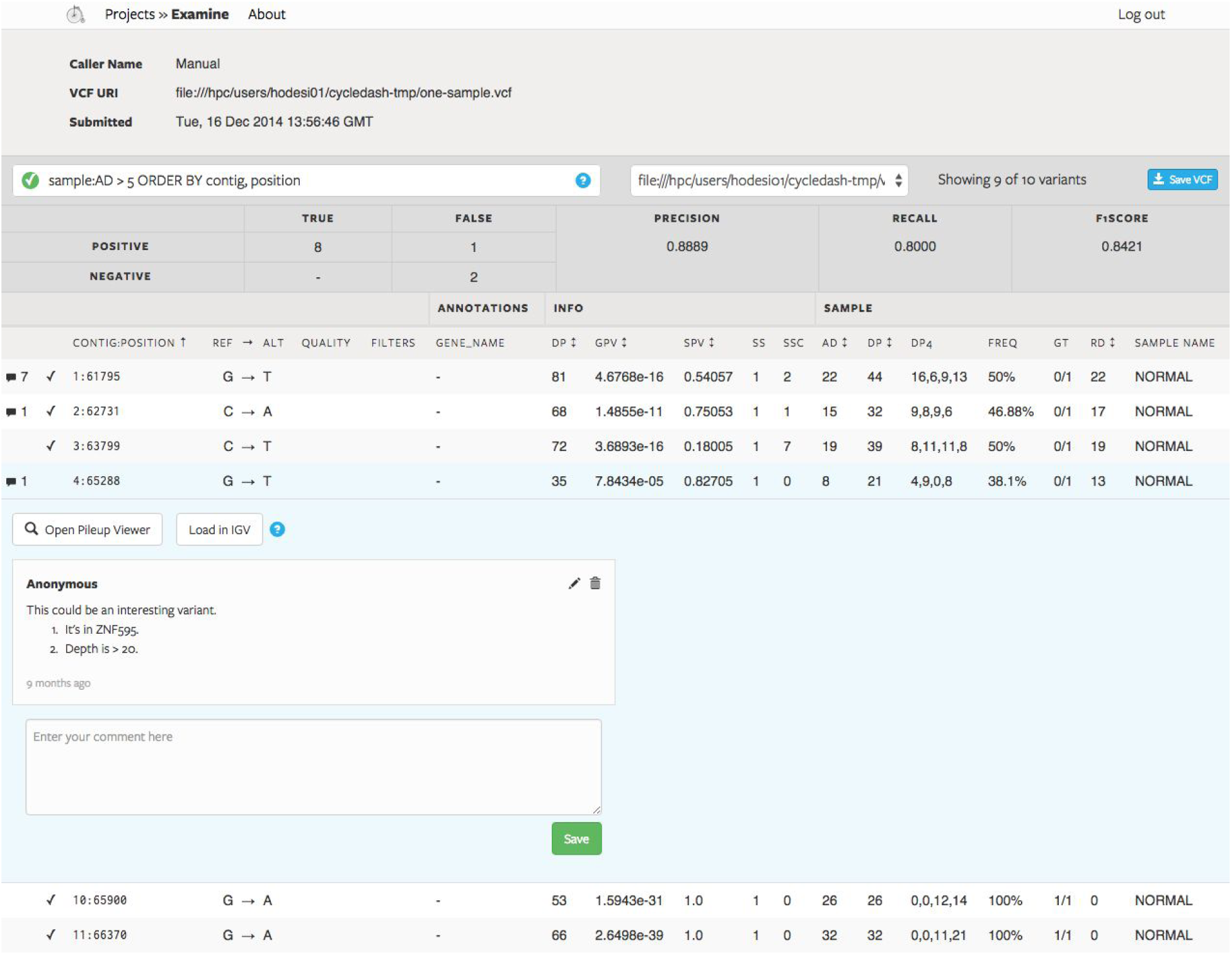
Cycledash “Examine” page

## Usage

Cycledash acts as a simple dashboard, accepting VCFs and BAMs via a RESTful interface, or from the web interface itself. We use a genomic workflow management tool, Ketrew (Mondet et al. 2016), to automatically post the results of our variant calling pipelines, which immediately begins the annotation and processing of the variants.

A user spends most of their time in the *examine interface* [fig 1], where they can filter variants by position, mutation type, any annotations that may have been added (such as gene name, or variant effect), and any other metadata found within the VCF, including read depth, variant allele frequency, and genotype. We provide a simple SQL-like interface for querying the variant table, implemented using PEG.js (https://pegjs.org/), allowing the user to sort as well as filter of variants. We also provide an interface for comparing variant collections against one another, so that concordance between variant callers, or a validation set, may be evaluated against the variant set. The output of any sorting and filtering may be exported to a VCF and downloaded from this page as well.

To facilitate collaborative analysis and the validation of somatic variants, we provide a per-variant commenting interface, as well as a simple way to “star” a variant, marking it as interesting and worth further exploration.

**Figure 2::**
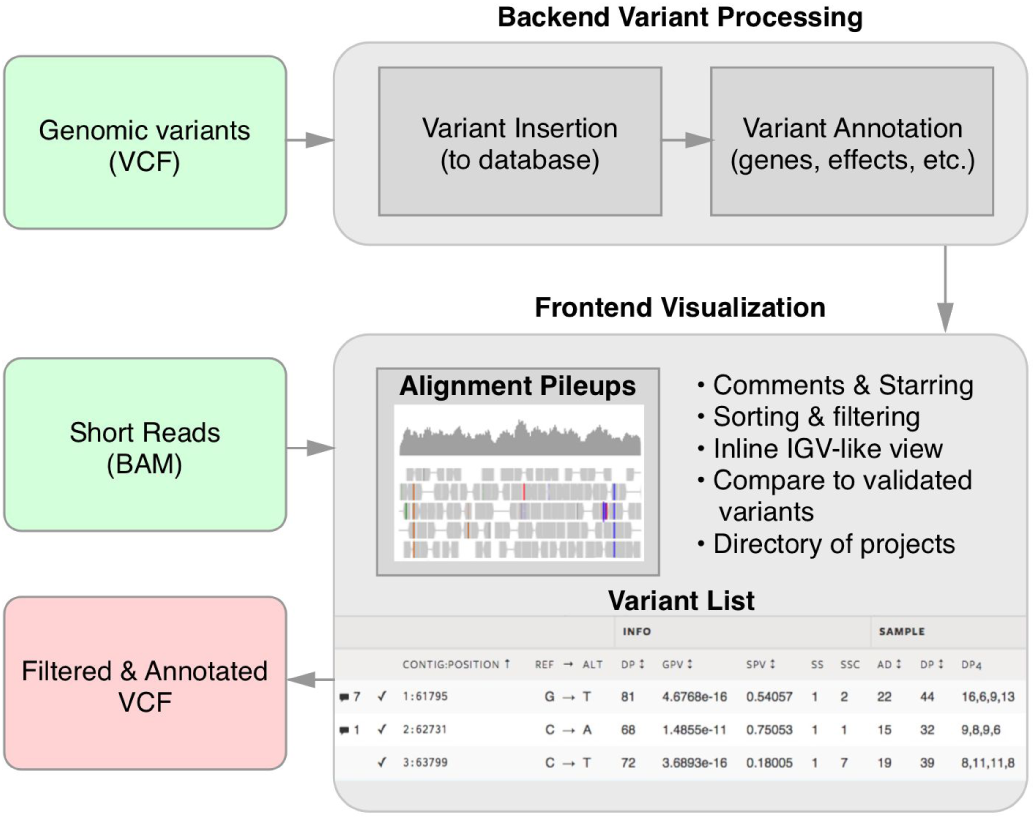
Flow of data through Cycledash.

Variants can be further inspected by opening the inline pileup viewer, Pileup.js. This allows the user to see the short read pileups in the area of the variant and determine if, for example, the variant caller may have made an error. This saves the user from having to leave Cycledash, locate the requisite BAM files, and open a pileup viewer.

## Acknowledgements

We thank all members of the Hammer Lab for their invaluable input on the manuscript and the application.

